# Genomic Language Models (gLMs) decode bacterial genomes for improved gene prediction and translation initiation site identification

**DOI:** 10.1101/2025.03.20.644312

**Authors:** Genereux Akotenou, Achraf El Allali

## Abstract

Accurate bacterial gene prediction is essential for understanding microbial functions and advancing biotechnology. Traditional methods based on sequence homology and statistical models often struggle with complex genetic variations and novel sequences due to their limited ability to interpret the “language of genes.” To overcome these challenges, we explore Genomic Language Models (gLMs) —inspired by Large Language Models in Natural Language Processing— to enhance bacterial gene prediction. These models learn patterns and contextual dependencies within genetic sequences, similar to how LLMs process human language. We employ transformers, specifically DNABERT, for bacterial gene prediction using a two-stage framework: first, identifying Coding Sequence (CDS) regions, and then refining predictions by identifying the correct Translation Initiation Sites (TIS). DNABERT is fine-tuned on a curated set of NCBI complete bacterial genomes using a k-mer tokenizer for sequence processing. Our results show that GeneLM significantly improves gene prediction accuracy. Compared to Prodigal, a leading prokaryotic gene finder, GeneLM reduces missed CDS predictions while increasing matched annotations. More notably, our TIS predictions surpass traditional methods when tested against experimentally verified sites. GeneLM demonstrates the power of gLMs in decoding genetic information, achieving state-of-the-art performance in bacterial genome analysis. This advancement highlights the potential of language models to revolutionize genome annotation, outperforming conventional tools and enabling more precise genetic insights.

## 1 INTRODUCTION

Bacterial gene prediction has been a long-standing focus in computational genomics, with numerous tools developed to tackle this challenge. Although significant progress has been made, gene annotation remains an evolving problem, particularly in the accurate identification of coding sequences and translation initiation sites. Traditional gene prediction methods such as Prodigal [1], Glimmer [2], and GeneMark [3] have been widely used for bacterial genome annotation. These tools rely on statistical models, heuristic-based rules, and sequence homology to infer gene structures, but limitations persist. One of the key issues in genome annotation is the variability in composition and organization. High-GC genomes, for example, pose unique challenges due to an increased number of potential open reading frames (ORFs) and ambiguous start codon selection, leading to a decline in prediction accuracy. Furthermore, annotation of Translation Initiation Site remains a difficult task, as start codon selection is influenced by diverse regulatory mechanisms that vary across species. Although existing tools such as TiCO [4] and TriTISA [5] have been developed to refine TIS predictions, these approaches miss several TIS predictions when tested using experimentally verified datasets. Another major challenge lies in the balance between sensitivity and specificity in gene prediction. Traditional methods often overpredict, frequently flagging numerous short ORFs that lack experimental validation. Moreover, genome sequencing analyses have revealed that many essential genes identified through transposon sequencing approaches contain false positives due to genome deletions [6], highlighting the need for methods that improve precision without sacrificing real gene annotations.

Recent advances in Artificial Intelligence and Machine Learning have revolutionized sequence analysis by enabling models to extract meaningful features from vast genomic datasets. Several deep learning-based models have been proposed for gene prediction, including convolutional neural networks (CNNs) [7, 8]. Self-supervised learning, a technique successfully applied in Natural Language Processing, has demonstrated remarkable capability in understanding sequential patterns and dependencies. Inspired by this progress, Genomic Language Models (gLMs) have emerged as a promising solution, treating DNA sequences as structured linguistic data to enhance gene annotation [9]. Using deep learning architectures, these models can go beyond traditional sequence alignment, capturing both local and global relationships within DNA sequences. Among these architectures, transformers, particularly Bidirectional Encoder Representations from Transformers (BERT) [10] have shown significant promise in sequence-based tasks. Unlike conventional gene prediction models, which rely on predefined feature sets, BERT-based gLMs dynamically learn sequence representations through self-attention mechanisms [11]. This capability allows them to infer gene structures with greater accuracy by modeling long-range dependencies within bacterial genomes. The application of such models offers a path toward more adaptive and precise genome annotation.

To address these challenges, we introduce a transformer-based genomic language model suited for bacterial gene prediction. We use a BERT-based architecture to tackle sequence classification tasks in Prokaryotes gene prediction. To address these tasks, we go through a two-stage classification process. As illustrated in Figure 1, given a genome, we have many ORFs that can be extracted. Among those ORFs, some of them cover non-coding regions, while others correspond to coding sequences. In the first step, we classify the coding sequences. Once this is done, in the second step, we focus on the TIS within the coding region. Specifically,

**Figure 1.**
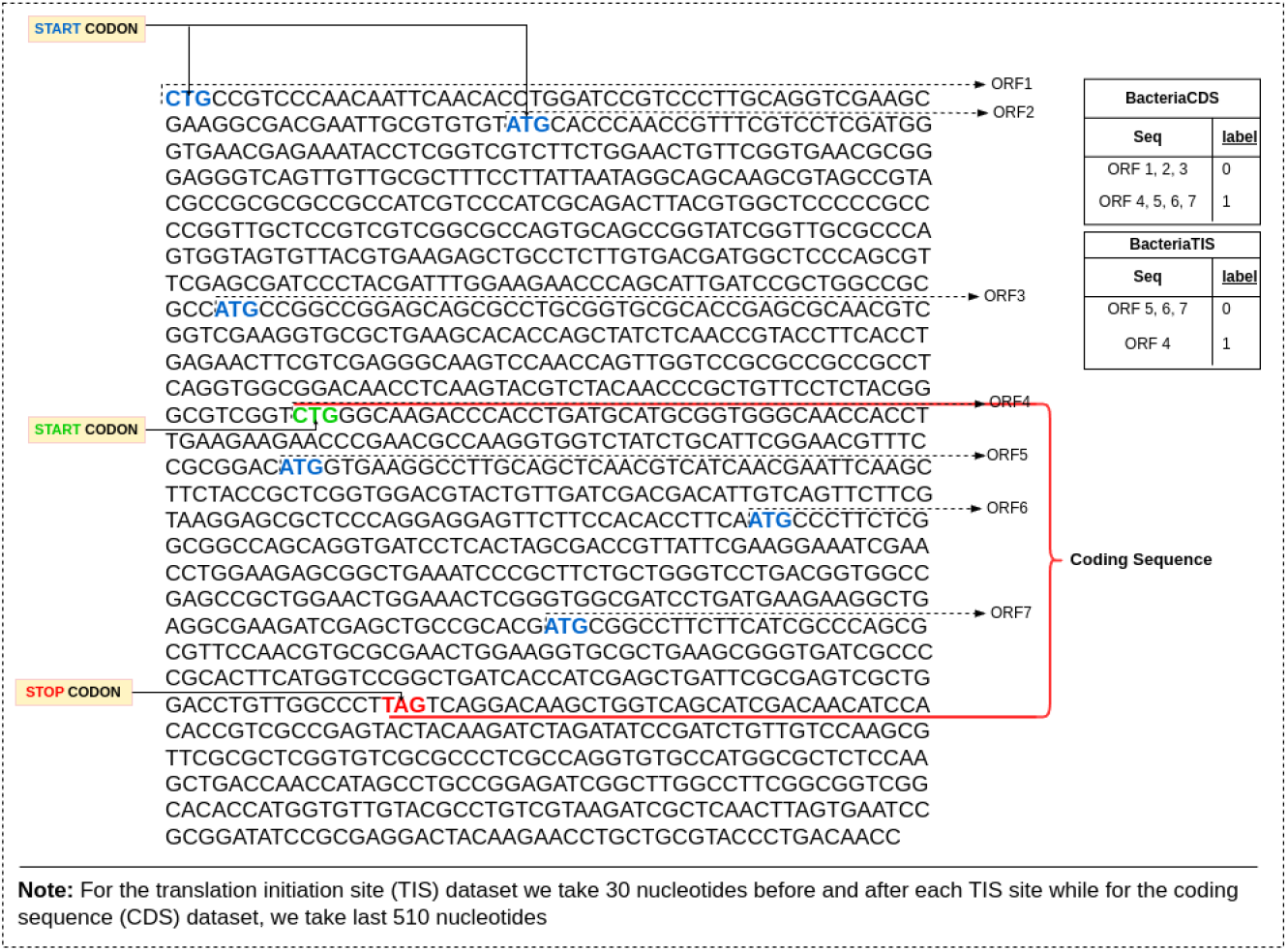
Illustration of Translation Initiation Site and Coding Sequence zones in the genome.

- A BERT-based genomic language model is developed to identify Coding Sequence regions and classify Translation Initiation Sites in Prokaryotes genomes.
- The model is initially trained using self-supervised learning in human genomic datasets to learn DNA fragment representations. It is then adapted for gene annotation through a two-stage pipeline: first, a model classifies ORFs into CDS and non-CDS regions, followed by a refinement stage where a second model identifies true TIS sites.
- Comparative evaluations are conducted against traditional gene prediction tools, assessing improvements in precision, recall, and scalability. A case study on verified bacterial genomes is performed to demonstrate the applicability and robustness of our framework in real-world genomic annotation tasks.
- An analysis is performed to interpret the decision-making process of the model, providing insight into its reasoning, an essential aspect of genomic research.

## 2 MATERIALS AND METHODS

### 2.1 Data collection

To develop a robust and generalized model, it is essential to curate a comprehensive and high-quality dataset. In this study, bacterial genomic data were obtained from the NCBI archive via the GenBank database. Specifically, the assembly summary file was downloaded from the NCBI at ftp.ncbi.nih.gov/genomes/genbank/bacteria. This file, approximately 1GB in size, provides extensive information on bacterial genomes, including annotations, assembly details, and metadata. To ensure dataset quality, only genomes with complete assembly status were considered and filtering was applied to retain only those classified under the “reference genome” category. This resulted in a dataset comprising `5,745 complete and annotated genomes`, representing `4,823 unique organisms`. For each genome in the filtered dataset, two essential files were retrieved: the `genome.fna` file, which contains the complete nucleotide sequences in FASTA format, and the `annotation.gff` file, which provides detailed gene annotations, including coding sequence information. The GFF file delineates genomic features, offering structured annotation data crucial for downstream processing and model training.

### 2.2 Data Processing

To construct high-quality datasets for the translation initiation site and coding sequence classification tasks, we developed a multi-stage processing pipeline. This pipeline extracts open reading frames from bacterial genomes, assigns labels based on genomic annotations and balances the dataset to ensure robust model training. To extract ORFs, we scanned the forward and reverse strands of each genome sequence using a slightly modified version of ORFipy [12], a fast and flexible Python-based tool for ORF extraction, to identify potential ORFs based on specific start and stop codons. We filter ORFs that began with one of the start codons (`ATG`, `TTG`, `GTG`, or `CTG`) and terminated at a stop codon (`TAA`, `TAG`, or `TGA`). Nested overlapping ORFs were retained to provide comprehensive genomic coverage. Each extracted ORF was subsequently assigned a label by comparing its genomic coordinates with the CDS annotations in the corresponding GFF file. We then labeled two types of datasets, one for CDS and the other for TIS classification. The CDS dataset comprises CSV files containing nucleotide sequences with a maximum length of 510, each labeled positive (1) or negative. A positive label indicates that the sequence represents the longest truncated ORFs whose start or end positions align with an annotated CDS in the GFF reference file. On the other hand, the TIS dataset includes only sequences from ORFs that match a CDS sequence in the reference file and sequences are assigned a binary label, where 1 represents a true translation initiation site (TIS). To capture sequence context, each TIS-centered sequence includes 30 nucleotides upstream and downstream, resulting in a total length of 60 nucleotides. The overall annotation process of both datasets is illustrated in Figure 2.

**Figure 2.**
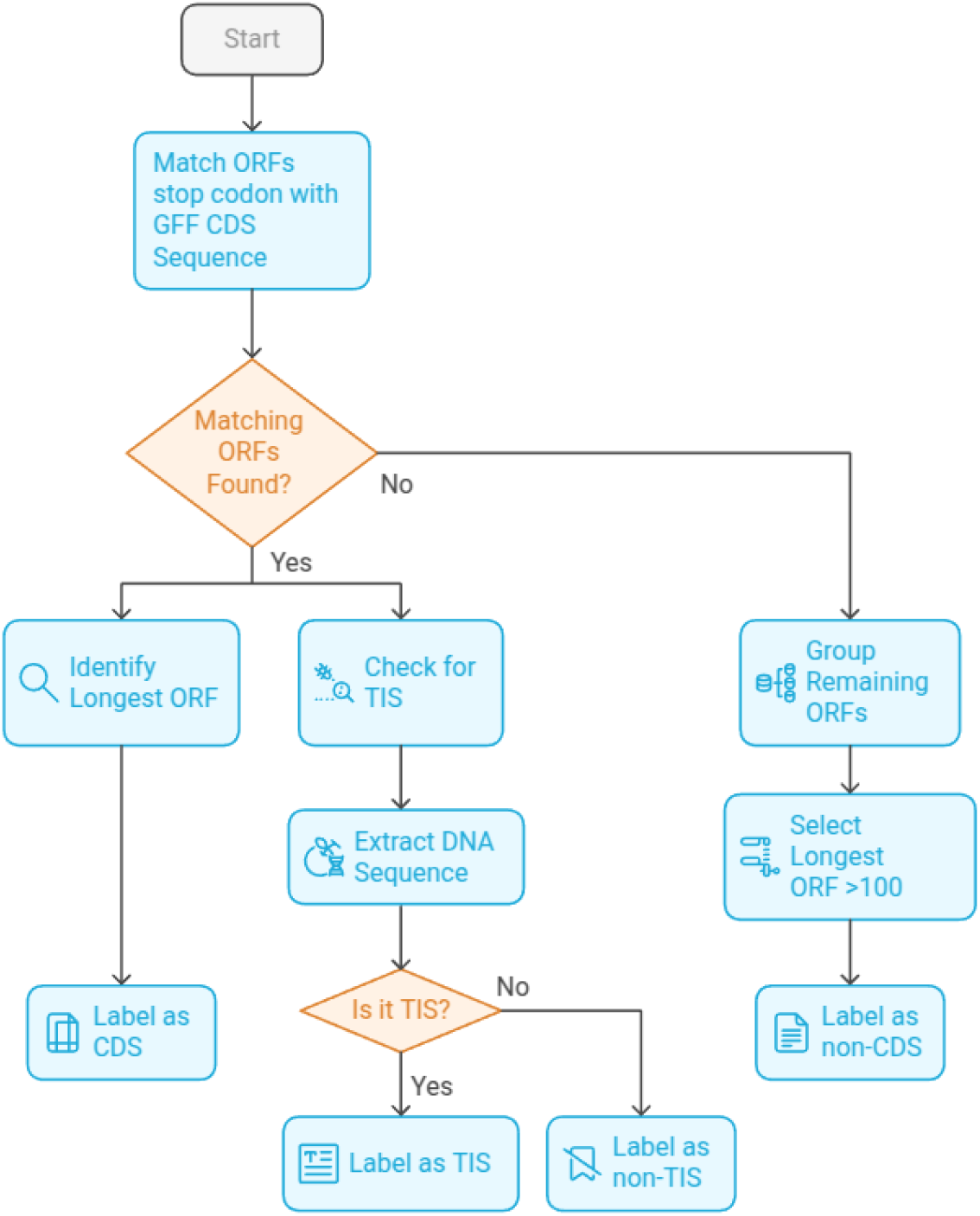
Overview of the dataset labeling pipeline for CDS and TIS classification. The pipeline involves extracting nucleotide sequences, assigning binary labels based on alignment with annotated CDS regions in the GFF reference file, and structuring the sequences for downstream analysis.

To ensure class balance and mitigate potential biases during model training, different sampling strategies were applied to the CDS and TIS datasets. For the CDS dataset, negative samples were downsampled based on sequence length to match the distribution of the positive class. This approach increases the difficulty of classification, encouraging the model to learn discriminative features beyond sequence length for distinguishing CDS from non-CDS regions. In contrast, for the TIS dataset, where all sequences have the same fixed length, random undersampling was performed to achieve class balance without introducing additional biases. The dataset was partitioned into training, testing, and evaluation sets. For the CDS dataset, the class-balanced splits consist of `14,975,672` sequences for training, `2,181,188` for testing, and `4,253,562` for evaluation. Similarly, for the TIS dataset, the partitioning resulted in `14,544,028` sequences for training, `2,199,192` for testing, and `4,136,340` for evaluation.

### 2.3 Tokenization & Embeddings

To process DNA sequences effectively during training, we employ the `k-mer encoding` technique used in the DNABERT [13] framework. In this method, the DNA sequence is split into overlapping substrings of length *k*, known as *k-mers*. Each k-mer serves as a discrete token, like words in natural language processing. According to the DNABERT paper, state-of-the-art performance was achieved using *k* = 6. Therefore, for our experimentation, we use a *k* = 6 encoding scheme. Given a DNA sequence, the tokenizer splits it into overlapping 6-mer tokens with a stride of 3 for the CDS classification task, while for TIS classification, we use a default stride of 1. After tokenization, each k-mer is transformed into a numerical representation using the pre-trained DNABERT model. Each k-mer is mapped to a fixed 768-dimensional vector, corresponding to the hidden size of the BERT [10] architecture. Additionally, we incorporate special tokens: `[CLS]` at the beginning and `[EOS]` at the end of each sequence. The `[CLS]` token provides an additional contextual representation for the sequence as a whole. As a result, for the CDS classification task, each sequence is represented as a matrix of size (512, 768), while for the TIS classification, each sequence is represented as a matrix of size (62, 768). This transfer learning enables the k-mer representations in each sequence to capture meaningful and contextually enriched information. Using DNABERT embeddings, we ensure that the tokenized sequences provide a robust foundation for downstream fine-tuning.

### 2.4 Model

Our model is based on the DNABERT framework, which extends the Bidirectional Encoder Representations from Transformers to genomic sequences. DNABERT follows the pre-training and fine-tuning paradigm, where the model first learns general DNA sequence representations and is later fine-tuned for specific downstream tasks. It retains the same transformer-based architecture as BERT, consisting of 12 self-attention layers, 768 hidden dimensions, and 12 attention heads, as shown in Figure 3.

**Figure 3.**
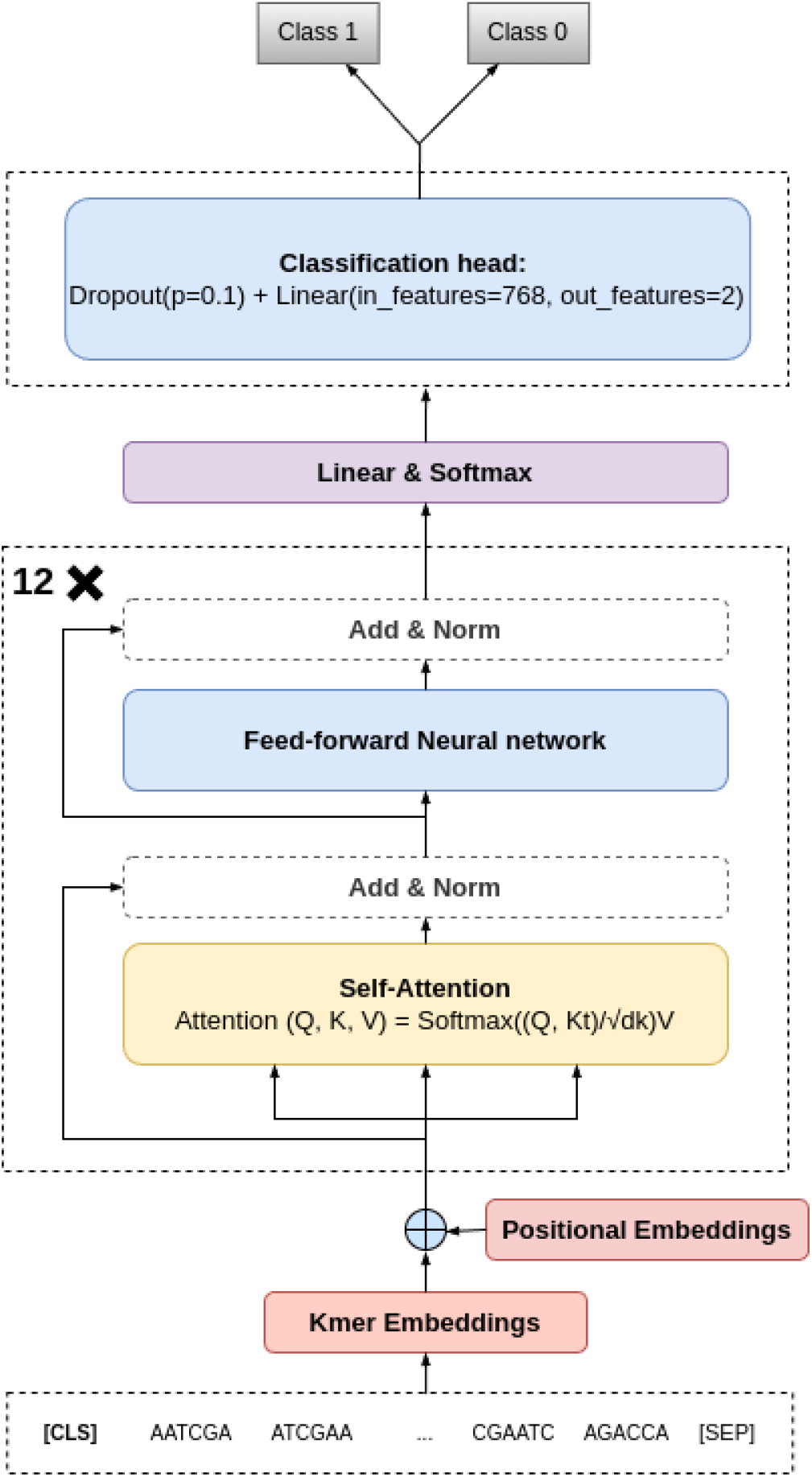
Illustration of the transformer-based BERT architecture used for genomic sequence classification. The model consists of multiple self-attention layers that process k-mer tokenized DNA sequences, capturing both local and long-range dependencies. The input sequence is embedded and passed through 12 transformer layers, each containing 12 attention heads and hidden dimensions of size 768. The output representations are fine-tuned for specific downstream tasks such as coding sequence classification and translation initiation site prediction.

Given a tokenized input sequence represented as k-mers, the model applies multi-head self-attention to learn contextual relationships across the sequence. The self-attention mechanism is formally defined as:

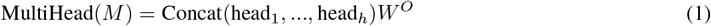

where each attention head is computed as:

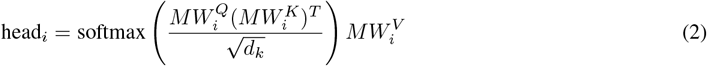

Here, *M* represents the input token embeddings, while 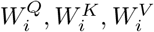 are the learnable query, key, and value matrices for the *i*-th attention head. The self-attention scores determine the contextual relevance of each token concerning all others in the sequence. The final hidden representations from the transformer layers are then used for sequence-level or token-level classification tasks.

### 2.5 Fine-Tuning

For a given sequence, we prepend the classification token `[CLS]` to the tokenized input sequence and then generate its embedding representation using the DNABERT base model. Self-attention is applied, where the query and key matrices are both derived from the sequence itself. As illustrated in Figure 4, the attention mechanism computes attention weights using a dot product between the query and key matrices. A softmax function normalizes these weights, and the result is multiplied by the value matrix to adjust the original input embeddings. The re-weighted embeddings are then passed through feed-forward layers, culminating in a classification head appended to the DNABERT base architecture. The final classification decision is based on the embedding of the `[CLS]` token, which aggregates global sequence features.

**Figure 4.**
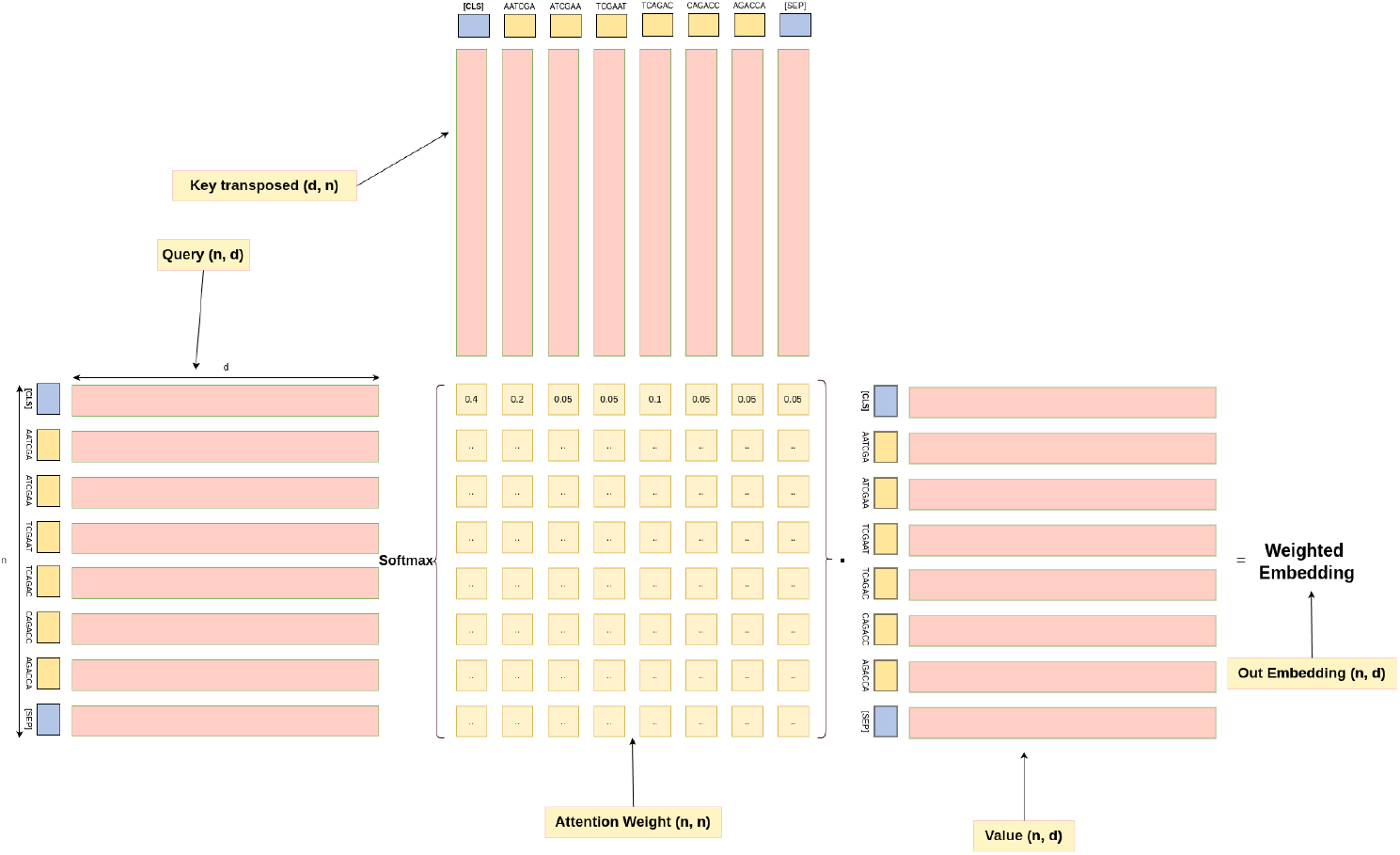
Self attention illustration: The attention mechanism assigns dynamic importance to different k-mer tokens within the sequence by computing attention scores from query, key, and value matrices. The scores are normalized using a softmax function, and the resulting weighted sum refines the sequence embeddings.

Our fine-tuning approach focuses on two key classification tasks: identifying Coding Sequence regions from DNA sequences and detecting Translation Initiation Sites within CDS regions. Fine-tuning follows the standard BERT-based classification procedure, where the final hidden representation of the `[CLS]` token is passed through a fully connected layer for sequence-level classification. To enable this, DNABERT is extended by appending a classification head, which consists of:

- **Pooler Layer**: Aggregates information from the `[CLS]` token embedding to generate a fixed-size representation for the entire sequence.
- **Fully Connected Layers**: A feed-forward neural network with one or more layers that maps the pooled representation to the output classes.
- **Softmax Layer**: Converts the output logits into probability scores for classification.

The objective function for fine-tuning is the cross-entropy loss, defined as:

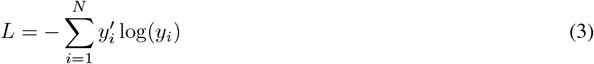

where 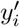 represents the ground-truth labels, and *y*_*i*_ denotes the predicted probabilities for each of *N* classes. In our case for both experiments *N* = 2.

### 2.6 Experiment Setup

The entire workflow, from input tokenization and embedding generation to final classification, is summarized in the pipeline diagram (Figure 5). This diagram outlines all key components, including the DNABERT model, Transformer layers, and the classification head. Given the large-scale dataset, we implement an iterable dataset using PyTorch’s data utilities to efficiently handle data loading and batch processing. The model is trained using the AdamW optimizer with a linear warm-up strategy. The learning rate is initialized at 3*e −*5 and gradually decayed to zero over the training steps. The model undergoes fine-tuning on NVIDIA GPUs partitions. To ensure robust evaluation, we partition the dataset into training and testing subsets, where the test set contains only organisms absent from the training set. This experimental design ensures that the model’s performance is assessed on previously unseen taxa, effectively evaluating its generalization ability.

**Figure 5.**
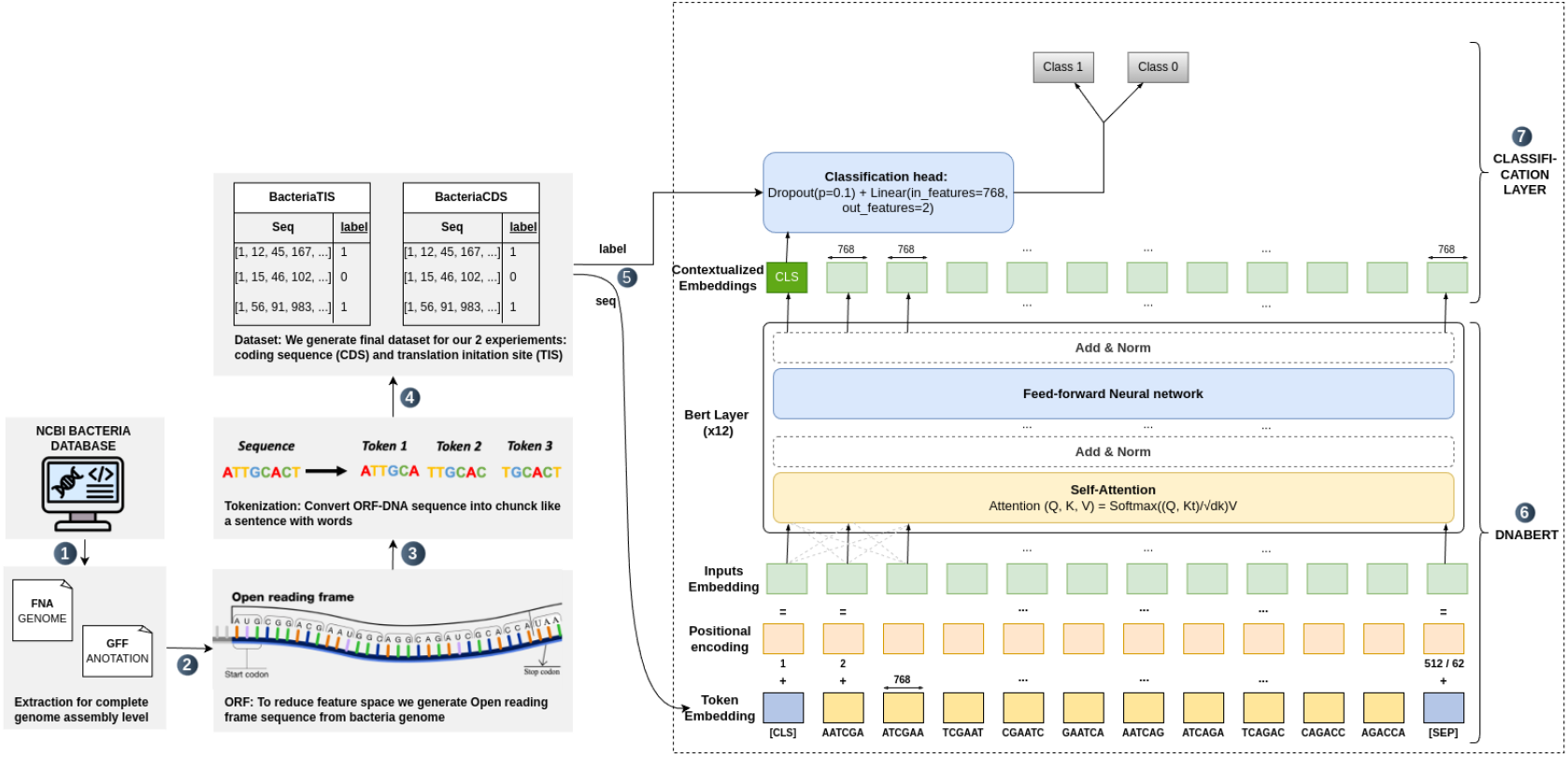
TIS & CDS fine-tuning pipeline: The procces involves extracting Open Reading Frames from bacterial genome data, tokenizing sequences into k-mers, and training a transformer-based model to classify genomic regions. The self-attention mechanism enables the model to capture long-range dependencies within DNA sequences, enhancing prediction accuracy.

#### CDS Classification

The objective of this task is to classify coding sequences in genomic data. The CDS classification experiments were conducted on the Toubkal supercomputer. The training was performed on two NVIDIA A100 GPUs, each equipped with 80GB of memory. Distributed training was implemented across both GPUs with a batch size of 512, and the total training duration was approximately 37 hours for 2 epochs.

#### TIS Classification

For Translation Initiation Site classification, the objective is to classify 60-bp sequences centered around the potential translation initiation site. This experiment was conducted at the [College of Computing, Bioinformatics Lab](https://bioinformatics.um6p.ma/) using a single NVIDIA RTX A5000 GPU with 24GB of memory. The model was trained with a batch size of 768, and the training process completed in approximately 19 hours for 3 epochs.

### 2.7 Evaluation Metrics

We used a set of standard classification metrics to evaluate the performance of both the individual binary classifiers and the final classifier (via stacking or max-voting). These metrics provide a comprehensive overview of the model’s ability to correctly classify protein transcription factor families. The following metrics were calculated:

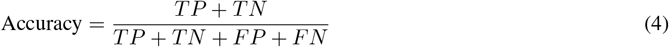

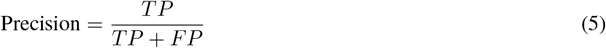

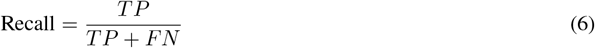

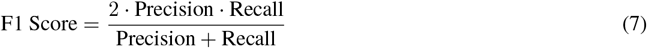

### 2.8 Post-processing

To facilitate seamless gene annotation after model training, we developed a post-processing pipeline that integrates an interactive web interface and an API-based system. The tool is publicly available on GitHub at Bioinformatics-UM6P/GeneLM. We implemented a web-based annotation tool that enables users to submit genome sequences for automatic annotation. The tool supports two modes of input: direct input where users can paste a genome sequence into the provided text area or file upload where users can upload a FASTA file for processing. Once the input is provided, users can specify the desired output format, selecting either GFF or CSV. Upon submission, the system processes the annotation and generates structured output files. The user-friendly interface ensures accessibility for researchers and bioinformaticians. A snapshot of the interface is shown in Figure 6.

**Figure 6.**
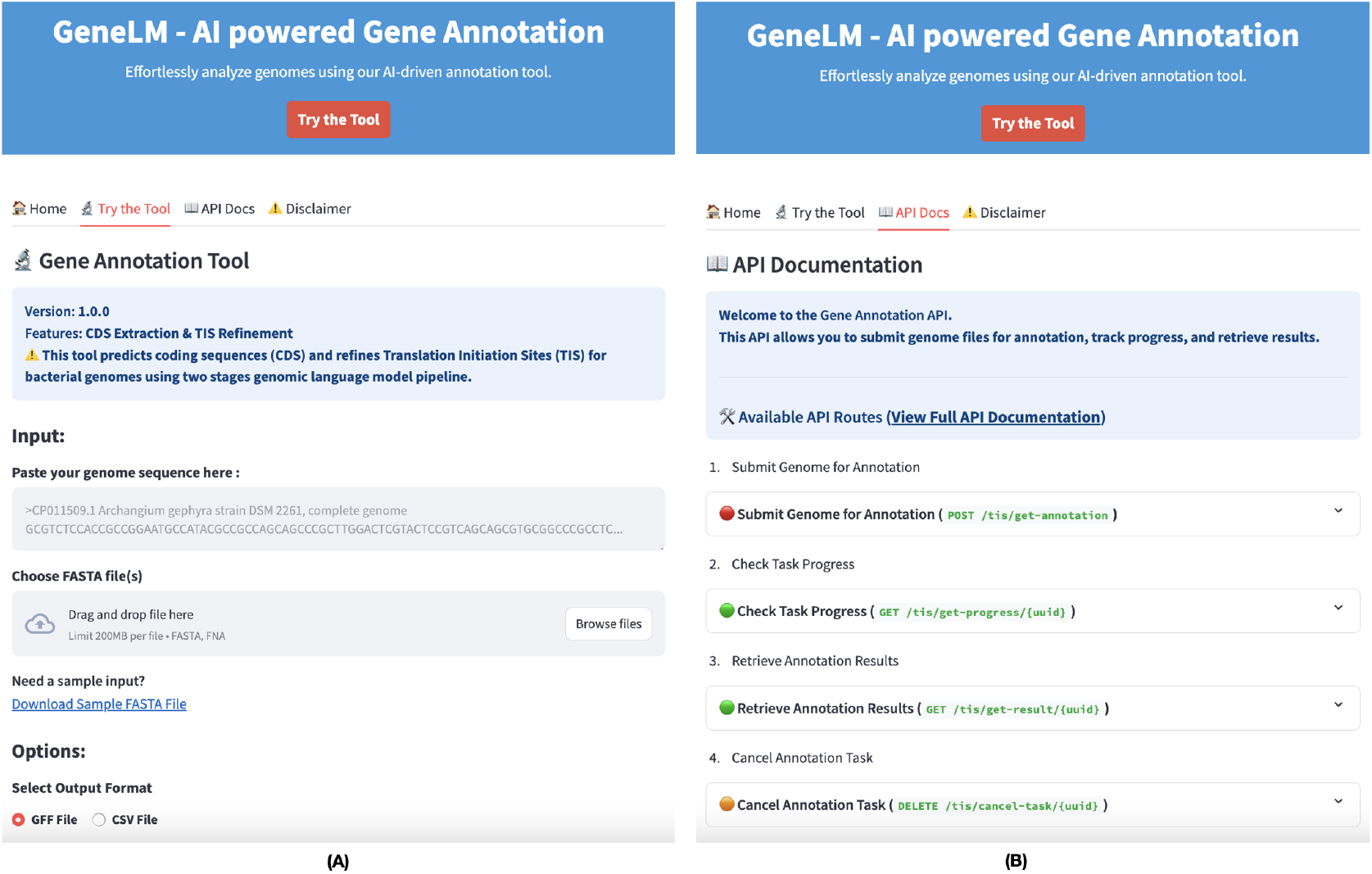
Web interface for genome sequence annotation. (A) The Gene Annotation Tool interface allows users to input genome sequences by pasting them or uploading FASTA files. It provides CDS extraction and TIS refinement functionalities with options to select the output format (GFF or CSV). (B) The API Documentation interface outlines available API routes for genome annotation, including submitting sequences, tracking progress, retrieving results, and canceling tasks.

Beyond the graphical interface, we developed a RESTful API to enable programmatic access for flexible genome annotation. The API allows users to submit annotation tasks asynchronously, queue multiple annotation jobs efficiently, track annotation progress, retrieve annotated results and cancel annotation tasks if necessary. The API is well-documented, providing multiple endpoints for annotation submission, result retrieval, and task management. This ensures seamless integration into bioinformatics pipelines and allows users to perform genome annotation directly through Python scripts or other computational workflows. An overview of the API’s functionality is depicted in Figure 6. This integrated web-based and API-driven annotation pipeline provides an efficient, scalable, and user-friendly solution for genome annotation, bridging the gap between machine learning-based gene prediction and real-world biological research.

## 3 RESULTS AND DISCUSSION

This section presents the results obtained from our experiments, structured into three key evaluation stages. First, we report the performance of our model during training and testing, assessing its effectiveness using standard evaluation metrics. Next, we evaluate the model’s performance in a real-world setting by testing it on experimentally verified sequences, providing a practical assessment of its predictive accuracy on biological data validated in the laboratory. Finally, we compare our approach against state-of-the-art gene annotation tools, benchmarking its accuracy and efficiency. These evaluations provide a comprehensive understanding of the model’s capabilities and its potential for practical applications in genome annotation.

### 3.1 Training performances

In this section, we present the results obtained from training our model across different experiments.

#### CDS Classification

Figure 7 illustrates the training loss progression over multiple steps and the comparison of evaluation metrics across two fine-tuning epochs. Initially, the training loss decreases sharply as the model learns, stabilizing after several thousand iterations. However, sporadic spikes in loss indicate instances of challenging gradient updates, possibly due to the model encountering difficult samples or high-weight adjustments. Despite these fluctuations, the overall trend shows a consistent reduction in loss, affirming effective learning. After the second fine-tuning epoch, the evaluation metrics showed marginal improvements (Table 1). The small gain in performance, combined with the computational cost of additional training, led us to halt fine-tuning at epoch 2. Table 1 presents the evaluation metrics for both training epochs and the final test set. The model achieved high classification performance, with precision, recall, and accuracy exceeding 98%. A slight reduction in evaluation loss suggests stable and effective training across epochs. The final model was evaluated on an independent test set, maintaining high accuracy (99.43%) while preserving a low evaluation loss (0.0211). These results confirm the model’s robustness and generalization capabilities.

**Table 1:** Evaluation and test set performance metrics for CDS classification.

| Metric | Evaluation Set |  | Test Set |
| --- | --- | --- | --- |
|  | Epoch 1 | Epoch 2 | Final Score |
| loss | 0.021233 | 0.020184 | 0.021132 |
| Precision | 0.981846 | 0.983868 | 0.983253 |
| F1 | 0.984364 | 0.985109 | 0.984544 |
| Accuracy | 0.994265 | 0.994553 | 0.994364 |
| Recall | 0.986916 | 0.986357 | 0.985844 |

**Figure 7.**
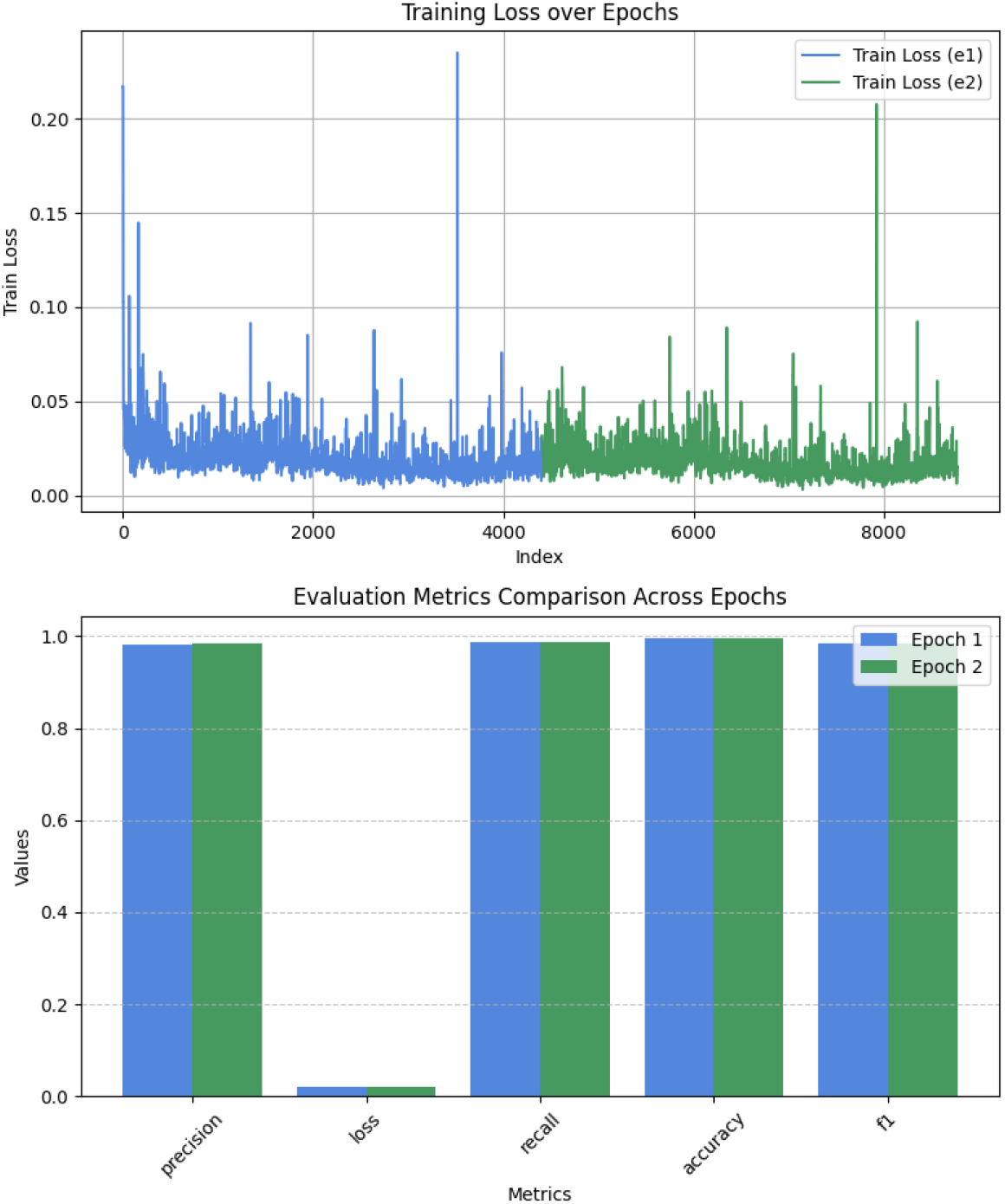
Train and Evaluation Metrics Comparison Across Epochs for CDS classification experiment.

#### TIS Classification

Figure 8 illustrates the training loss trend and evaluation metrics for the TIS classification experiment. Similar to CDS classification, the training loss shows a steep initial decline before stabilizing, with minor oscillations. The evaluation metrics in Table 2 show consistent performance improvements across epochs, with precision, recall, and F1-score increasing slightly over each iteration. The final test evaluation confirmed an accuracy (94.13%) and an evaluation loss (0.1546). One should note that the TIS information available on NCBI and used to compute the above accuracy is done using automatic gene prediction tools which may not always reflect the ground truth. To further validate our results, we compare our tool with existing tools using all known experimentally verified TIS data available for bacterial genomes.

**Table 2:**
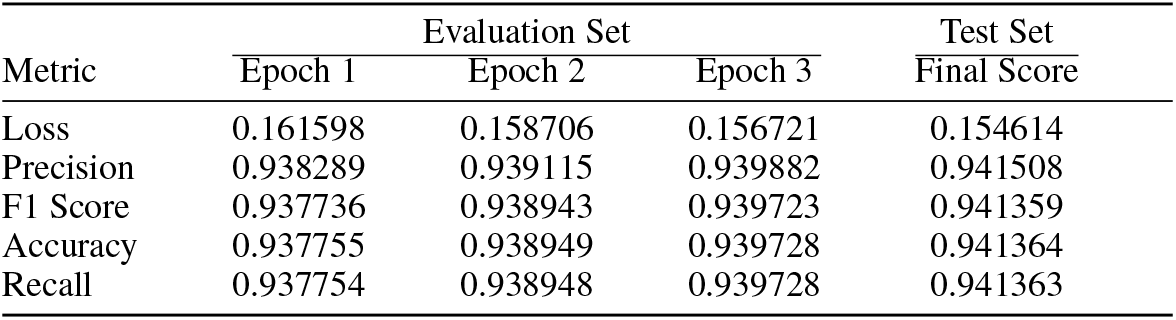
Evaluation and test set performance metrics for TIS classification for both training epochs and the final test set.

**Figure 8.**
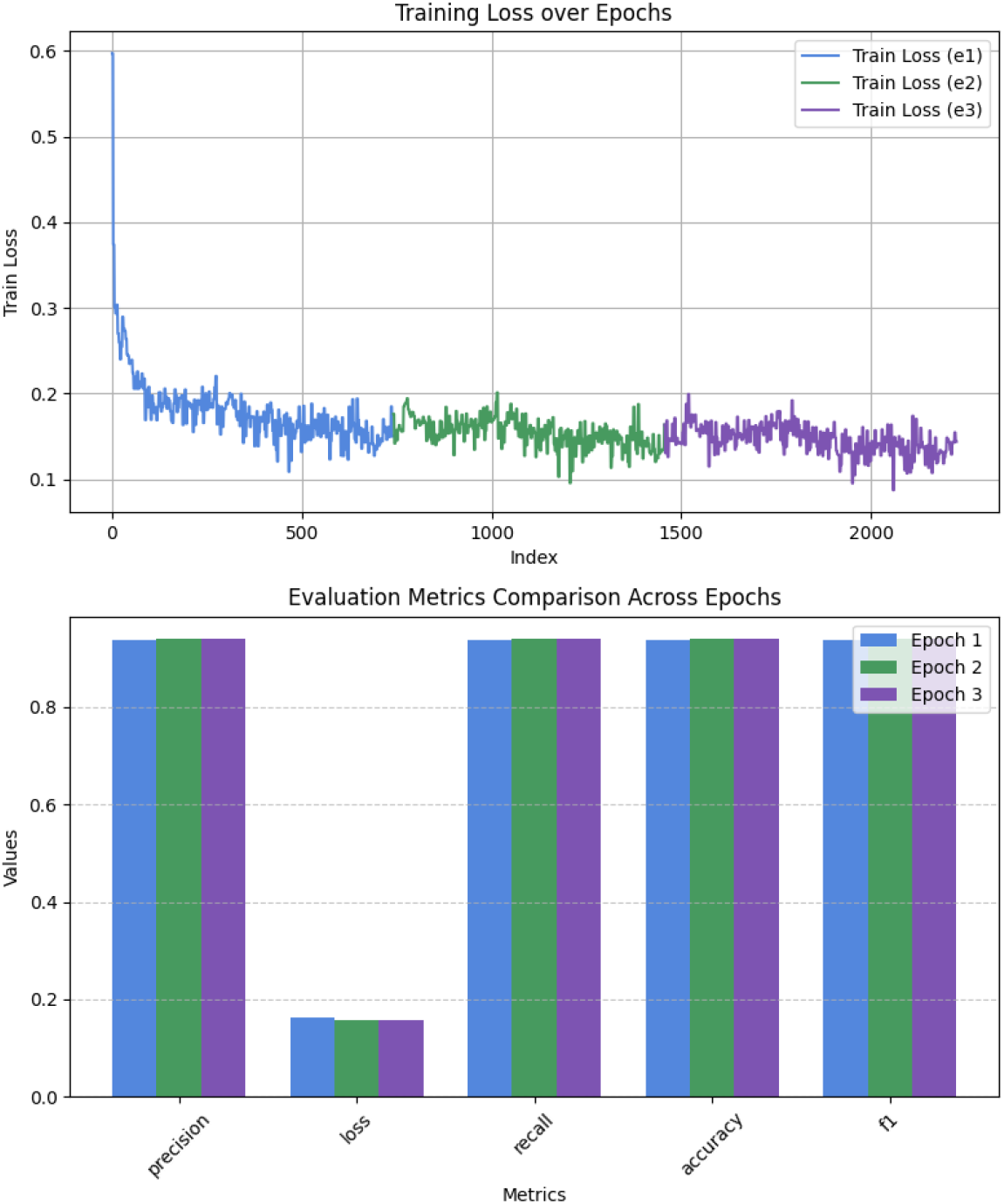
Train and Evaluation Metrics Comparison Across Epochs for TIS classification experiment.

### 3.2 Performance Benchmarking on Experimentally Verified Genomes

The effectiveness of GeneLM was benchmarked against Prodigal [1] using experimentally verified coding sequences (CDS). Benchmarking with real biological data is a standard practice for evaluating gene prediction methods, as seen in prior studies such as TRITISA [5]. To ensure real-world applicability, we retrieved verified reference sequences for Escherichia coli, Halobacterium salinarum, Natronomonas pharaonis, Mycobacterium tuberculosis, and Roseobacter denitrificans. The FNA files of these bacteria were processed using our TIS prediction pipeline (GeneLM) and also analyzed with Prodigal v3.0.0-rc.1, installed from its official GitHub repository. We compared the matched TIS, missed TIS, and total found CDS by both methods, using the total verified TIS as the reference. Our results indicate that while Prodigal remains a state-of-the-art tool, our approach provides superior performance in accurately identifying Translation Initiation Sites. Table 3 summarizes the key findings. As illustrated by this table, the results indicate that our approach consistently detects more verified TIS sites compared to Prodigal, with significantly fewer missed predictions. Notably, for E. coli K-12, Prodigal missed 431 sites, while our approach reduced this number to only 25. Similar trends were observed across all tested organisms.

**Table 3:** Benchmarking results comparing Prodigal and GeneLM for experimentally verified TIS.

| Bacteria | Total Verified TIS | Prodigal |  |  | GeneLM |  |  |
| --- | --- | --- | --- | --- | --- | --- | --- |
|  |  | Matched TIS | Missed TIS | Total Found CDS | Matched TIS | Missed TIS | Total Found CDS |
| *Escherichia coli* K-12 MG1655 | 769 | 338 | 431 | 4347 | 744 | 25 | 4213 |
| *Halobacterium salinarum* R1 | 530 | 243 | 287 | 2851 | 438 | 92 | 2659 |
| *Mycobacterium tuberculosis* H37Rv | 701 | 311 | 390 | 4204 | 626 | 75 | 3853 |
| *Natronomonas pharaonis* DSM 2160 | 315 | 169 | 146 | 2873 | 248 | 67 | 2737 |
| *Roseobacter denitrificans* Och114 | 526 | 0 | 526 | 4120 | 492 | 34 | 4006 |

### 3.3 Explainability: TIS Prediction motifs visualization

The task of Translation Initiation Site classification involves categorizing 60-nucleotide windows surrounding the TIS site. To understand the model’s decision-making process, we analyze how attention mechanisms contribute to classification. Figure 9 demonstrates that the model captures distinct patterns through its attention heads. For example, as observed in Figure 9, layer 1, head 11, focuses on a region approximately 30 base pairs upstream of the TIS, which may indicate the presence of a promoter site. In contrast, layer 2 exhibits a more selective focus on upstream regions, and a similar pattern is noticeable in layer 9.

**Figure 9.**
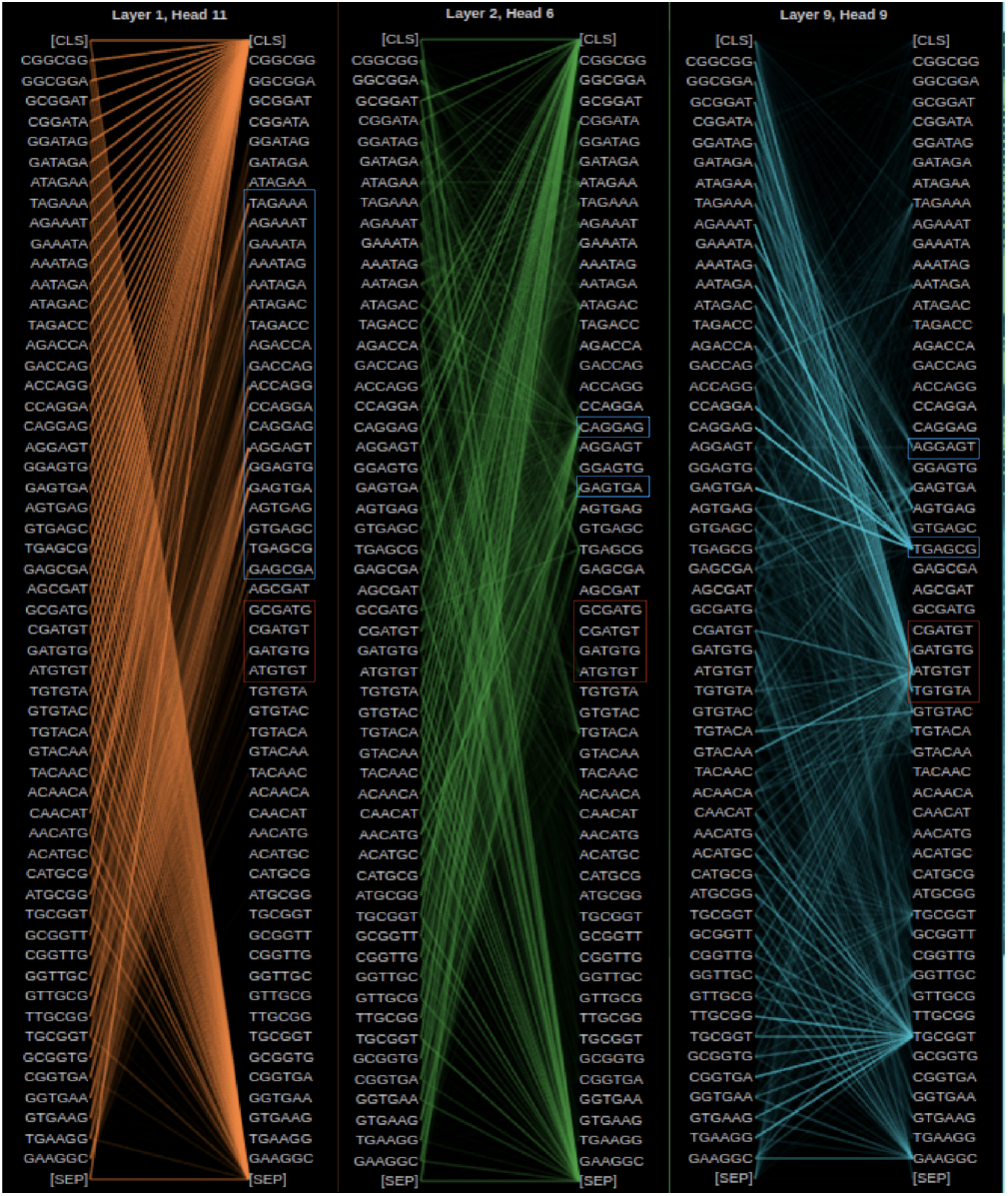
Visualization of attention mechanisms in the TIS classifier generated with BertViz tool [14].

To derive more generalized insights, instead of analyzing isolated sequences, we extend our visualization across all sequences in our verified test set, which consists of five bacterial species. We focus on layer 11, the final attention layer, as it aggregates all prior learned patterns and directly influences the classification head, playing a crucial role in the final decision. For true TIS sites, we compute attention weights using our fine-tuned TIS prediction model, yielding fixed-size tensors of 11 heads with dimensions 57 × 57 (computed as 60 - 5 + 2). The mean attention weights are calculated over all 11 heads per sequence and then averaged across all TIS instances.

Figure 10 presents the attention weight landscape for each bacterial species. The heatmaps labeled **(a), (b), (c), (d)**, and **(e)** correspond to *Escherichia coli, Halobacterium salinarum, Mycobacterium tuberculosis, Natronomonas pharaonis*, and *Roseobacter denitrificans*, respectively. The red box highlights the immediate region surrounding the TIS, while the blue box reveals an intriguing pattern: it marks sequence regions that receive high attention from the classification (CLS) token. These upstream regions may contain potential promoter elements, as the CLS token is responsible for sequence-level classification. This pattern is systematically observed across all bacterial species and aligns with the expected biological significance of promoter regions. Figure 11 further illustrates the discovered pattern between the classification token and upstream TIS positions for each bacterial species, emphasizing the model’s consistent focus on biologically relevant regions.

**Figure 10.**
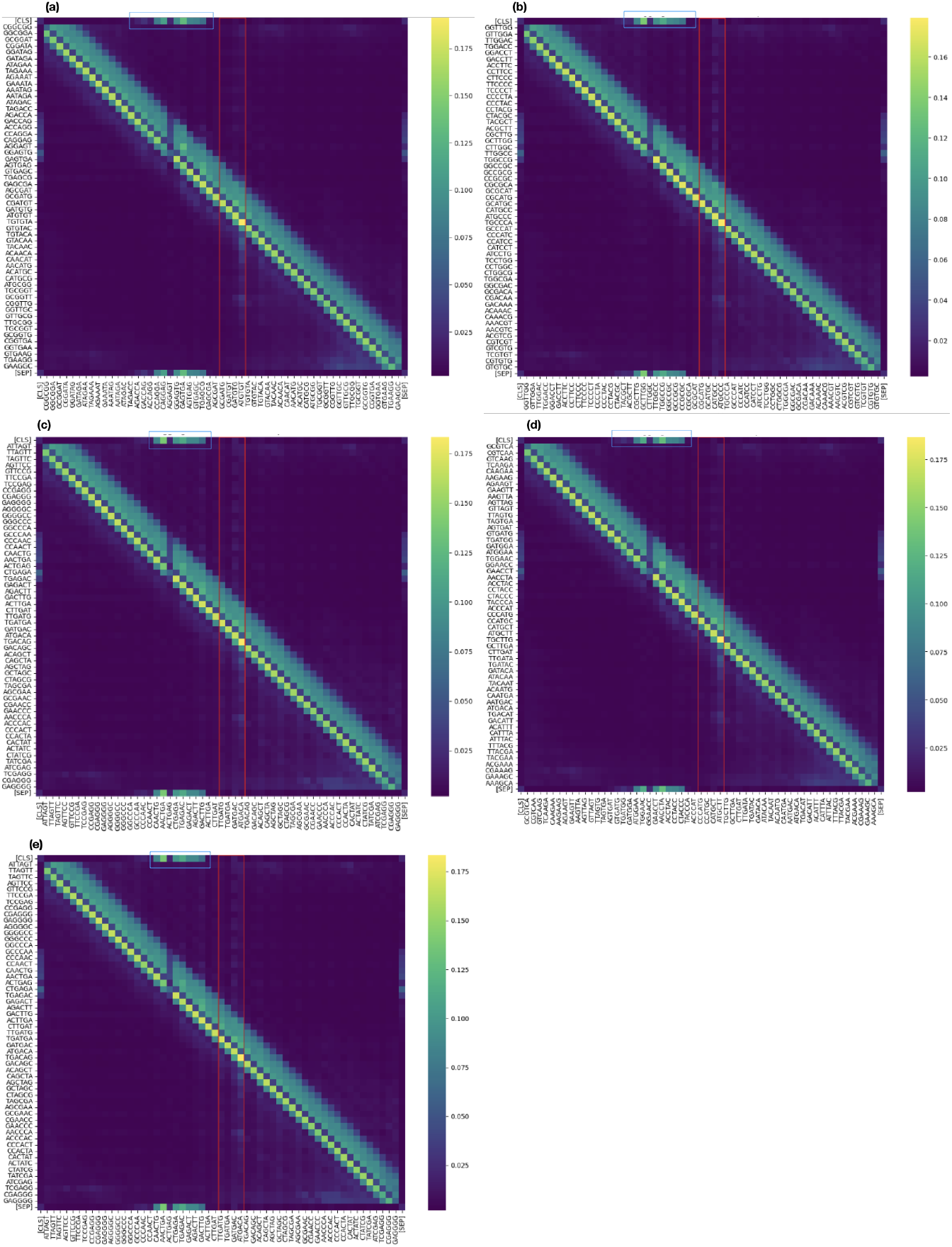
Mean attention weight distribution across different bacterial species. Each heatmap represents a verified bacterial species, where the mean attention weights are computed for sequences labeled as true TIS. This visualization highlights the regions where the classification token `[CLS]` focuses the most when making the final decision.

**Figure 11.**
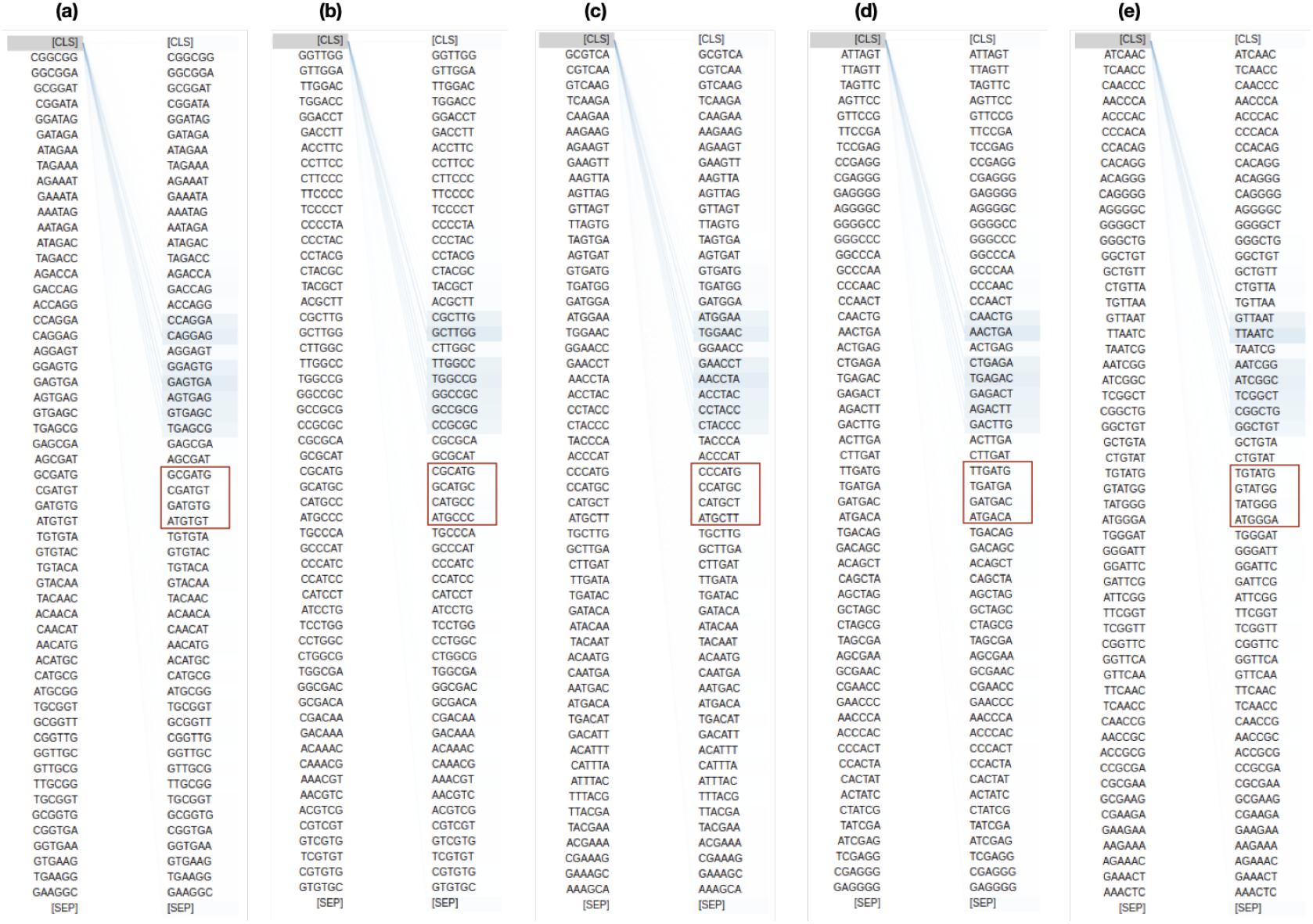
Observed systematic pattern between the CLS token and TIS upstream positions for a sample sequence.

## 4 CONCLUSION

The results obtained from our study highlight the effectiveness of transformer-based genomic language models for bacterial gene annotation. Our approach, based a BERT-based architecture, demonstrated significant improvements in both Coding Sequence classification and Translation Initiation Site detection compared to traditional gene annotation tools. One of the key findings of this study is the ability of the model to capture biologically relevant sequence patterns through self-attention mechanisms. The visualization of attention maps revealed that the classifier systematically focuses on upstream regions of TIS, potentially corresponding to promoter regions, across different bacterial species. This consistency suggests that the transformer-based model successfully identifies meaningful regulatory elements, reinforcing its applicability in genomic sequence analysis. Another crucial aspect of our findings is the generalization capability of the model. The high classification accuracy across diverse bacterial species underscores the robustness of the proposed approach. Additionally, the benchmarking results against traditional tools such as Prodigal indicate that our model significantly reduces false positives while maintaining high recall rates, ensuring better annotation precision without compromising sensitivity. Despite these promising results, some limitations remain. While transformer-based models provide improved sequence representations, their computational demands are higher compared to heuristic-based gene prediction tools. The requirement for large-scale training datasets and high-performance computing resources can be a barrier for widespread adoption in resource-constrained environments. Future work will focus on enhancing model interpretability and optimizing computational efficiency.

## Code and data availability

The current implementation of our work and the trained models and their weights are available at: Bioinformatics-UM6P/GeneLM.

## Acknowledgments

The authors express their gratitude to the College of Computing at Mohamed VI Polytechnic University for providing access to the supercomputing resources (https://cc.um6p.ma/toubkal-super-computer) used in conducting the research presented in this paper.

